# extgfa: a low-memory on-disk representation of genome graphs

**DOI:** 10.1101/2024.11.29.626045

**Authors:** Fawaz Dabbaghie

## Abstract

The representation of genomes and genomic sequences through graph structures has undergone a period of rapid development in recent years, particularly to accommodate the growing size of genome sequences that are being produced. Genome graphs have been employed extensively for a variety of purposes, including assembly, variance detection, visualization, alignment, and pangenomics. Many tools have been developed to work with and manipulate such graphs. However, the majority of these tools tend to load the complete graph into memory, which results in a significant burden even for relatively straightforward operations such as extracting subgraphs, or executing basic algorithms like breadth-first or depth-first search. In procedurally generated open-world games like *Minecraft*, it is not feasible to load the complete world into memory. Instead, a mechanism that keeps most of the world on disk and only loads parts when needed is necessary. Accordingly, the world is partitioned into chunks which are loaded or unloaded based on their distance from the player. Furthermore, to conserve memory, the system unloads chunks that are no longer in use based on the player’s movement direction, sending them back to the disk. In this paper, we investigate the potential of employing a similar mechanism on genome graphs. To this end, we have developed a proof-of-concept implementation, which we called “extgfa” (for external GFA). Our implementation applies a similar chunking mechanism to genome graphs, whereby only the necessary parts of the graphs are loaded and the rest stays on disk. We demonstrate that this proof-of-concept implementation improves the memory profile when running an algorithm such as BFS on a large graph, and is able to reduce the memory profile by more than one order of magnitude for certain BFS parameters.

**Availability:** Our implementation is written in `Python` and available on Github under the MIT license https://github.com/fawaz-dabbaghieh/extgfa

## 1. Introduction

Graphs and graph theory have been studied in discrete mathematics since Leonhard Euler’s paper on the Seven Bridges of Königsberg problem in 1736 (3). In computer science, the need for a versatile data structure that can represent data and its relationships led to the development of the graph data structure. Graph data structures have been well studied and have applications in a vast array of problems within and beyond the domain of computer science (10, 35). In the field of bioinformatics, researchers have been investigating the potential of graph data structures for the compact representation of sequencing data and the advantages this approach offers in the analysis of large-scale sequencing datasets. In a similar vein, numerous bioinformatics algorithms have been developed that rely on the use of graphs as a means of representing sequencing data. These graphs are typically referred to as “genome graphs” and have been employed to address a range of issues, including genome assembly (2, 34, 36), pangenome and proteome representation (24, 26, 31), sequence alignment (9, 23, 25, 33), visualization (5, 16, 38), and other related problems.

With this increased use of genome graphs, researchers faced new difficulties associated with storing, processing, and analyzing these graphs efficiently as the size of the graphs increased with more data. For example, recently, the Human Pangenome Reference Consortium (HPRC) published their draft human pangenome graph made of 47 phased and assembled diploid samples (26). Several graphs were produced depending on the technology used. For example, the graph produced using the Minigraph-Cactus method (19) has a raw file size of 48 Gb and contains 92,879,580 vertices, and the graph produced using the Pangenome Graph Builder pipeline (PGGB) (14) has a raw file size of 89 Gb and contains 110,884,673 vertices. To tackle this challenge, numerous software toolkits for working with large genome graphs were developed (13, 14). Furthermore, algorithms for searching, subgraph identification, and indexing of genome graphs were also developed (8, 20, 37). However, all these toolkits tend to load the entire graph in RAM (Random Access Memory) even if only a small part of the graph is needed.

As graph data structures have been studied in the computer science field for decades, the concept of using external or disk memory in lieu of RAM has been explored previously. This includes, for example, external memory breadth-first search (27), external memory depth-first search (18), and other external memory algorithms (1, 6). However, in these theoretical studies, researchers primarily focused on adapting a specific algorithm to allow external memory, but did not present a multipurpose external-memory graph data structure where any graph algorithm can then be implemented.

In this work, we present an idea and a proof-of-concept implementation for a general-purpose external memory representation of a graph in the GFA format (Graphical Fragment Assembly format) (15). This representation is inspired by open-world video games and how they manage memory usage. We demonstrate that even for a simple proof-of-concept implementation, we were able to reduce the memory profile by more than one order of magnitude for certain experiments and certain parameters. Moreover, we illustrate the limitations of our method and propose avenues for future improvements.

## 2. Methods

### 2.1. External memory in video games

In procedurally generated or open-world video games, storing the entire world in RAM is extremely inefficient and, for many games, simply infeasible. In such cases, the developers needed to come up with ways to keep small parts of the world in RAM, and have the ability to load more from disk seamlessly and without affecting the performance of the game (12, 32). One example of such games is *Minecraft* (28), a procedurally generated open-world video game, where the game’s world is built from different blocks consisting of different materials (sand, rock, grass, etc.). Moreover, the world is split into chunks and extends in each cardinal direction. To keep the gameplay as smooth and playable as possible, and to keep the RAM from overloading, chunks are stored on disk and only loaded into memory when the player is of a certain distance of the chunk (29).

Figure 1 shows a simple representation of the idea behind some of the video games like Minecraft. The green block in which the player is located represents the portion of the map that is fully loaded in memory. The yellow adjacent blocks represent parts of the map that are only partially loaded. For example, this could be distant features such as trees, mountains, houses, etc. that have not yet been fully populated with all aspects of the game, thus preserving the feeling of a vast open world while saving memory. Finally, the red blocks represent unloaded parts of the map.

**Fig. 1.**
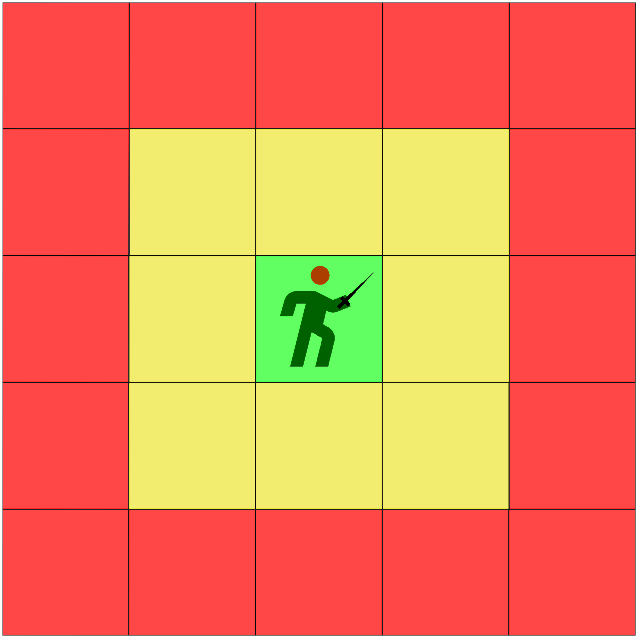
This figure shows a simple representation of how some open-world games like Mincraft manage their memory blocks. The green block is where the player is, and represents the part of the map that is fully loaded. The yellow blocks are parts of the map that are partially loaded, and the red blocks are parts that are not loaded.

When the player moves in a certain direction, the status of the map block changes according to the player’s movement direction, i.e., farther away parts or map chunks from the player become inactive and are removed from RAM, and other chunks in the direction of the player’s movement get loaded, which keeps the number of loaded map blocks in memory about the same.

### 2.2. Graph Chunking Pipeline

The main takeaway of memory management in open-world games is that the whole world does not need to be loaded into memory, only the map chunks that surround the player. Taking inspiration from this, we developed the following pipeline to chunk the GFA graph and produce an index that allows us fast access to parts of the graph instead of the complete graph, and allows us to keep loading and unloading parts of the graph as needed. The pipeline consists of the following steps:

## 1. Cutting the graph into neighborhoods

This step aims to cut the graph into non-overlapping chunks, where a chunk here is some connected subgraph or a community smaller than the original graph. Community detection in graphs is an old problem that has been explored for decades; many algorithms have already been developed to solve this problem with varying degrees of sensitivity, specificity, and time and memory complexity (22). More on the specific algorithms used in our tool can be found in the “Implementation” section.

## 2. Recursive chunking

Depending on the algorithm used to cut the graph into chunks, the chunks might not be balanced in terms of their number of nodes, and the sizes can vary tremendously. It is not strictly necessary for this method to have similarly sized chunks, however, this helps in keeping the loading and unloading time and memory of chunks uniform across the chunked graph. To achieve that, an upper and a lower threshold of the number of nodes per chunk can be used to limit the sizes of chunks. The upper threshold is the maximum number of nodes one chunk is allowed to have before getting cut further into smaller chunks; the lower threshold is the minimum number of nodes one chunk is allowed to have before getting merged - if possible - with a neighboring chunk. This step is run recursively until all chunks left have a size between the upper and lower thresholds.

## 3. Producing reordered GFA and indexes

After the previous steps, we are left with a set of non-overlapping chunks with sizes between the upper and the lower thresholds, where all the chunks jointly comprise the original graph. Using this information, we can reorder the GFA file, such that the nodes and edges of one chunk are written consecutively in the ordered GFA file, comprising one block, before starting with the next chunk. While writing the reordered GFA file, we keep track of the beginning and end positions of each chunk, allowing us later to load any arbitrary chunk from the file into RAM without having to read the complete GFA file. Moreover, a database relating each node ID to its corresponding chunk ID is also produced, to enable the retrieval of any given node in the graph.

## 3. Implementation

### 3.1. Graph Partitioning

Our implementation is written in `Python`, and uses the `NetworkX` library to run the community algorithms (17). Thereafter, it uses a custom GFA class for reading, writing, and manipulating GFA graphs. We call this proof-of-concept implementation `extgfa` (for External GFA).

For the graph cutting step, we tested several community algorithms already implemented in the `NetworkX` library, such as the Kernighan-Lin algorithm (21), edge betweenness partition (11), Louvian communities (4), and Clauset-Newman-Moore greedy modularity maximization algorithm (7). We found that the last algorithm works best compared to the others; it was relatively fast even for large graphs, and produced communities that are similar in size in terms of the number of nodes.

In brief, the Clauset-Newman-Moore greedy modularity maximization algorithm tries to find sets of nodes or “communities”, where each community is more densely connected internally than to other communities. This is achieved by starting with each node as its own community, then joining pairs of communities that maximize the “modularity”, until further merging does not increase the modularity. Modularity here can be simply explained as the score that maximizes the number of edges in a community compared to edges between communities (30).

We arbitrarily assign each produced chunk a unique integer ID starting from 1. From there, three files can now be generated that encapsulate the information of the chunks and allow dynamic loading and unloading of chunks into RAM when working with the chunked graph. Figure 2 shows an example graph represented in a GFA file format and visualized with `Bandage` (38). The three files produced by our implementation are:

**Fig. 2.**
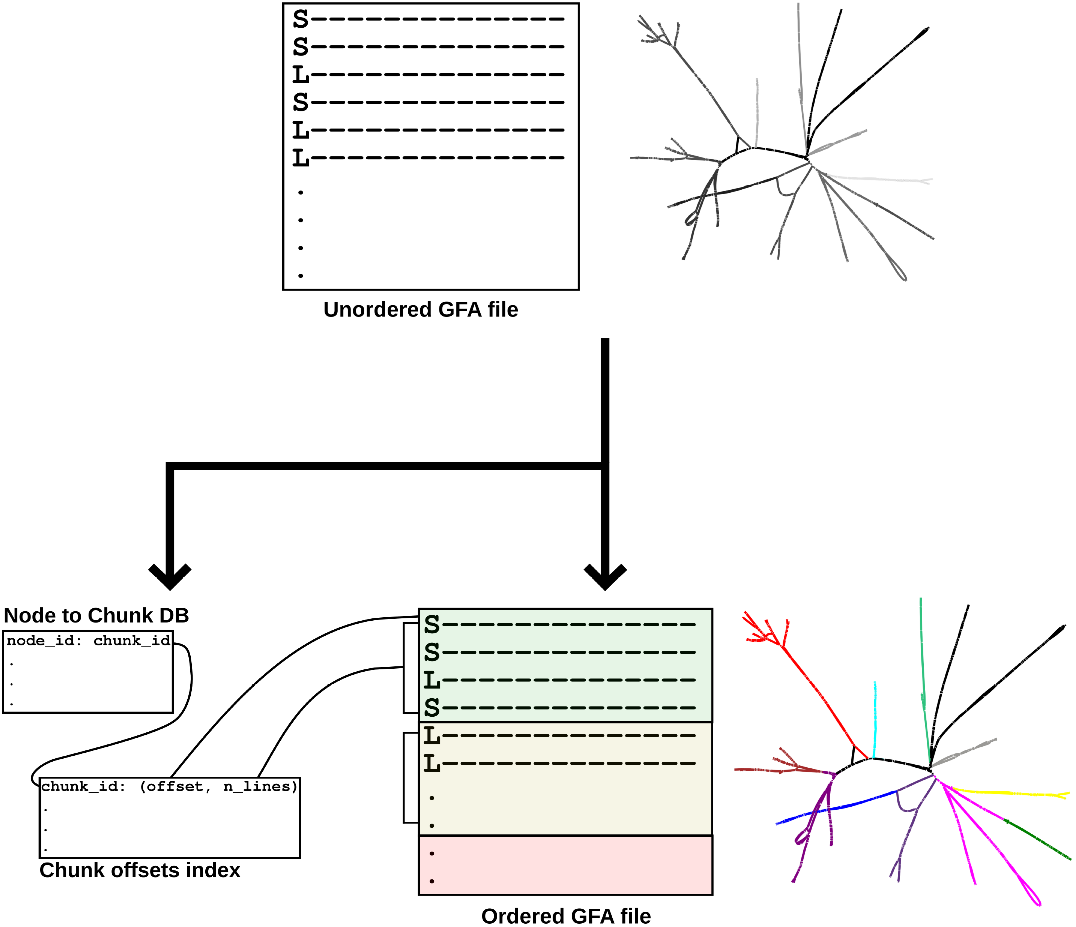
This figure represents the `extgfa` pipeline. First, chunks are detected in the GFA graph as mentioned in Section 2. Once we have the chunks, `extgfa` produces three files: a database `dbm` of key-value pairs where the keys are the node IDs and the values are the IDs of the chunk IDs the node belongs to; a binary file storing key-value pairs, the key being the chunk ID and the value a tuple of the file offset in the GFA and the number of lines to read from that offset; and a reordered GFA file where each chunk is written consecutively

1. A reordered GFA file, where chunks are written consecutively as blocks in the output file.
2. A chunk offset index, which consists of key-value pairs, where the key is the integer chunk ID and the value is a tuple of two values; the first value points to the offset in the reordered GFA file for that chunk, and the second value is the number of lines to read starting from the offset.
3. A `dbm` file built with the `shelve` library in `Python`, that builds a database of key-value pairs stored on disk. A `dbm` is a library or a database with single hashed keys that point to some value and provide fast access to the data stored. Values can be retrieved from this database without having to load it completely in RAM. In this database, the key is the node’s string ID and the value is the integer chunk ID.

The idea here is that for any node in the graph, we can retrieve the ID of the chunk it belongs to, then use that to get the reordered GFA file offset (i.e., where to start reading), and the number of lines to read for the entire chunk.

### 3.2. Chunked Graph Class

We implemented two similar graph classes, `Class::Graph` and `Class::ChGraph`, where the former loads the GFA graph completely, i.e., stores all the nodes and edges in memory, while the latter makes use of the three files produced previously to dynamically load and unload chunks as needed. Both classes have the exact same internal functions and data structures which allows a direct comparison between the two classes.

With the `Class::ChGraph`, it does not need to load any nodes or chunks at the beginning, only once the user tries to retrieve a node, the class then retrieves the chunk ID associated with that node using the node ID-chunk ID database, then finds the offset and number of lines to read in the reordered GFA file, and finally retrieves the chunk that the node belongs to. Moreover, the user can set a cutoff on how many chunks are allowed to be loaded in memory before older chunks are removed. This is done using a FIFO (First-In-First-Out) queue that keeps track of the chunks loaded. Once the threshold is crossed, the chunks loaded first are removed and new chunks are loaded and added to the queue. This queue is used to mimic how some video games unload the chunks that are further away from the player as the player moves.

Both classes implement basic functionality related to graphs, such as finding edges, nodes, node contents (sequence, length, tags, etc), graph traversals, and other functionalities.

However, `Class::ChGraph` is able to automatically recognize when it needs to load a new chunk without the user’s intervention. Therefore, a user can implement the same algorithm and use either of the classes without having to interact with any chunk loading or unloading functionality directly, which allows for a seamless integration.

## 4. Results

Here, we used the graph representing Chr22 from the HPRC PGGB V1 graph (14). The graph contained 3,759,736 nodes and 5,224,421 edges. To cut the graph into chunks, we used the Clauset-Newman-Moore algorithm as mentioned in the “Implementation” section, and we set the upper threshold to |*V*| */*2000 and the lower threshold to |*V*| */*5000, where |*V*| is the number of nodes in the graph. This resulted in 3,084 chunks with an average chunk size of 1,219 nodes per chunk. To test both graph classes, we implemented a standard BFS (Breadth-First Search) algorithm that starts from a user-defined node and traverses the graph until it either hits a user-provided size cutoff (BFS size) or finds no more nodes to traverse.

We chose a random starting node in the graph and ran the BFS algorithm with different BFS cutoff sizes (50, 100, 1,000, 5,000, 10,000, 50,000, 100,000, 500,000, and 1,000,000). Additionally, for the chunked version, we ran each BFS cutoff size with 7 different chunks queue sizes (1, 5, 10, 50, 100, 500, 1,000).

Figure 3 shows a scatter plot comparing running the BFS algorithm on both classes with the different BFS cutoff sizes; the top plot shows the memory profile and the bottom one shows the time profile.

**Fig. 3.**
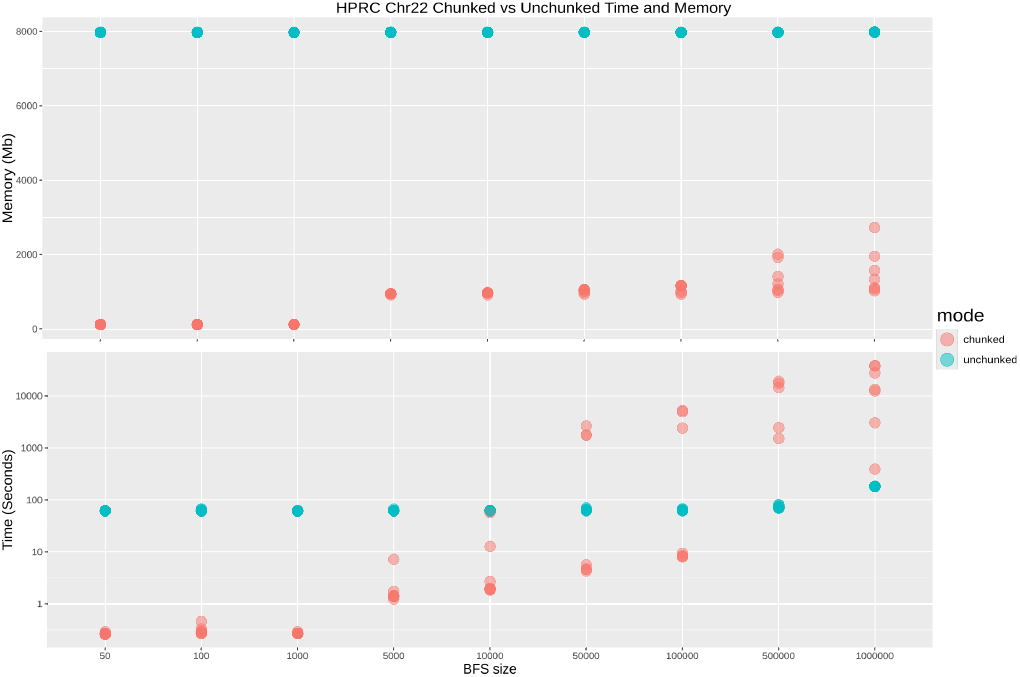
Scatter plot comparing both chunked and unchunked versions in terms of time and memory. We see that for the unchunked version, the time and memory are mostly constant, because we always need to load the complete graph, and this operation takes much more time compared to running the BFS algorithm. In contrast, we see more variability in terms of time and memory in the chunked version, which can be explained by the number of chunk load and unload operations needed, and the effect of the maximum number of chunks allowed in memory.

As the top plot shows, the unchunked version (loading graph completely) has a constant memory profile of approximately 8 Gb for all the BFS cutoff sizes. This is expected, as the unchunked version loads the complete graph before running the algorithm, which would always result in the same memory profile regardless of the BFS size. In contrast, for the chunked graph, the memory profile is smaller and affected by the BFS cutoff size, with a maximum memory usage of approximately 3 Gb. This is also expected, as the smaller the BFS cutoff size, the fewer chunks needed to be loaded into memory.

In the bottom plot of Figure 3, showcasing the time profile, the unchunked version behaves similarly to the top plot, with an approximately constant time profile (approximately 90 seconds), with a slight elevation in time for BFS cutoff size of 1,000,000, which is due to the time needed to perform the BFS algorithm. For the chunked version, we see a great variability in the time profile, especially for bigger BFS cutoff sizes. This has two reasons: The first reason, the bigger the BFS size, the more chunk-loading operations need to be performed, including database lookups and finding chunk offsets in the GFA file. The second reason is the maximum threshold of chunks in memory, which explains more the variability in time for a certain BFS size.

As mentioned in the “Implementation” section, once the chunk queue is full, older chunks need to be unloaded from memory. Because the loading and unloading is not happening on a separate thread, this will have a big effect on the processing time. Therefore, in order to explore the effect of the size of the chunk queue on both time and memory, and explain the variability in time profile of the chunked version, we plotted the individual values for each run of BFS on the chunked graph version only. The points are colored from light blue to dark blue based on the smallest queue size to the biggest; the results are shown in Figure 4. We see that for smaller BFS cutoff sizes, the effect of the chunks queue size becomes negligible, which can be attributed to the fact that for smaller BFS cutoff sizes, we only need to load one or very few chunks into memory. However, the effect grows as the cutoff is larger. We can see the expected memory-vs-time trade-off: as more chunks can be loaded into memory at a time, more nodes are readily available for fast access, avoiding slowing down the BFS algorithm and increasing the memory requirement, and *vice versa*.

**Fig. 4.**
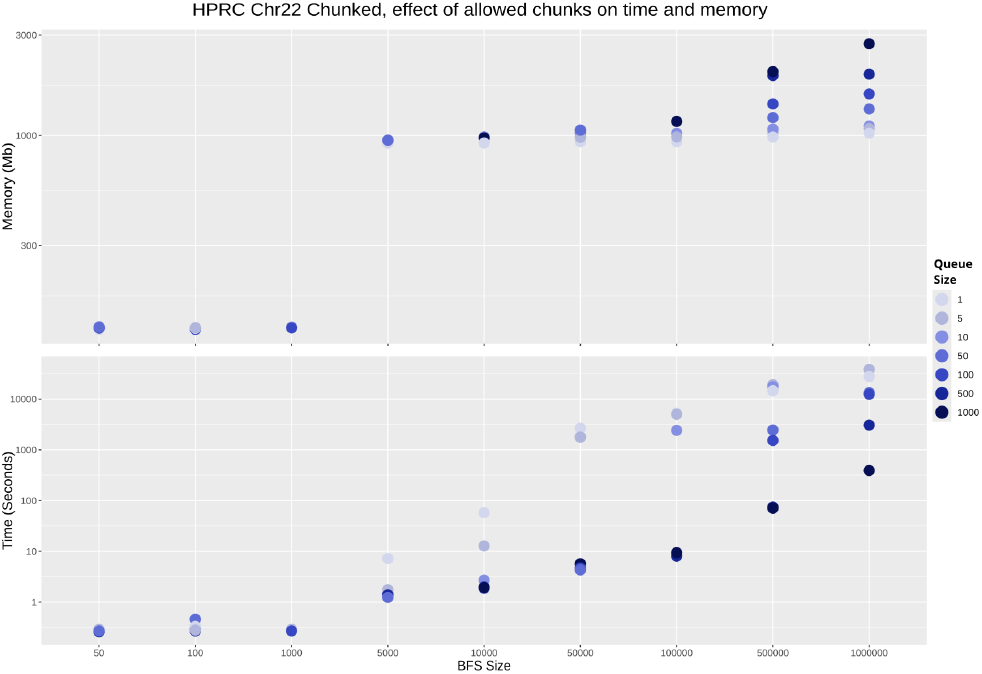
Scatter plot showcasing the effect of the chunks queue size on both memory and time in the chunked graph version. We see that the bigger the BFS cutoff size is, the greater the effect the queue size has. Moreover, the queue size has a contrasting effect on time and memory; the bigger the queue, the less time needed to run the BFS and the more memory required, and *vise versa*

## 5. Discussion and Conclusion

In this paper we presented `extgfa`, a proof-of-concept `Python` implementation for an external-memory representation of genome graphs in GFA format. We showcased that the memory profile could be improved by partitioning the graph into smaller chunks stored on disk, with a mechanism for fast access and retrieval of these chunks only when needed. Moreover, for certain use cases and parameters, the time profile also improved considerably when compared to loading the complete graph in memory first.

We believe that with the fast development of graph genomes in bioinformatics, and the increasing shift towards pangenomes instead of linear reference genomes, a dynamic on-disk data structure for working with such graphs is needed. Furthermore, solutions which stem from other computer-science and related fields also dealing with large amounts of data that cannot be stored in memory can be applied to effectively solve this problem.

Even though our implementation is labeled as a proof-of-concept implementation, it is usable, especially for cases where the user would like to investigate many subgraphs or loci in a large graph and does not want or is unable to load the complete graph into memory. `extgfa` can then be used to extract a size-bounded subgraph around a user-specified start node.

Further improvements on the design and implementation can, of course, be made. For example, similar to how some video games load chunks around the player, and keep loading in the direction the player is moving, before the player steps on these parts, one could implement chunk prefetching in such a way as to load neighboring chunks concurrently while certain nodes are being explored. Another possibility is to reduce memory usage by loading only the topological structure without loading any of the node’s other information such as sequence.

## ACKNOWLEDGEMENTS

We would like to thank Tobias Marschall for his input and continuous support, and much appreciation goes to Konstantinn Bonnet for many fruitful discussions and encouragements that resulted in this work. We would also like to thank Rebecca Serra Mari and John Stavrellis for looking over the manuscript and providing valuable input.

